# Expert-Guided Supervised Annotation of Erythroid Differentiation in Single-Cell RNA-seq

**DOI:** 10.64898/2026.06.05.730313

**Authors:** Alessio Enderti, Nicolò Stranieri, Simone G. Riva, Jim R. Hughes

## Abstract

Accurate annotation of intermediate cell states remains a major challenge in single-cell RNA sequencing (scRNA-seq), particularly in continuous differentiation systems such as erythropoiesis. Existing reference-based methods often lack the resolution required to distinguish early and transitional erythroid progenitors and may generalise poorly across datasets and modalities. Here, we present a supervised framework for erythroid lineage annotation based on expert-curated training data that integrates bulk and single-cell transcriptomic information. Starting from a human bone marrow scRNA-seq atlas, we refined erythroid annotations by introducing previously unresolved progenitor stages, including burst-forming unit–erythroid (BFU-E), colony-forming unit–erythroid (CFU-E), and pro-erythroblast (ProE), guided by canonical marker genes and bulk RNA-seq references. We trained and benchmarked four classical machine learning models and identified LightGBM as the best-performing approach, achieving a validation macro F1-score of 0.821 and balanced accuracy of 0.826. On a held-out test set, the model showed strong performance across most erythroid stages, with errors largely confined to adjacent differentiation states. The classifier was further transferred to independent bulk RNA-seq samples and an external bone marrow scRNA-seq dataset, where it recovered expected erythroid progression and refined coarse-grained annotations into higher-resolution cell states. Together, these results show that expert-curated supervised learning can improve erythroid cell state annotation in scRNA-seq and provide a practical framework for studying differentiation hierarchies in settings where finely resolved public references are limited.

## I Introduction & Motivation

Single-cell RNA sequencing (scRNA-seq) has transformed the study of cellular heterogeneity and differentiation in complex tissues [1], [2]. By profiling transcriptomes at single-cell resolution, it enables the identification of rare populations, the reconstruction of continuous developmental trajectories, and the inference of lineage relationships inaccessible to bulk approaches [3], [4]. These advances have been especially impactful in haematopoiesis, where cell fate decisions unfold along structured yet dynamic differentiation hierarchies.

A central challenge in scRNA-seq analysis is the accurate annotation of cell types and intermediate states. Unsupervised clustering can identify transcriptionally distinct populations [5], [6], but assigning biologically meaningful labels to these clusters remains a bottleneck. Manual annotation depends on marker gene inspection and domain expertise, introducing subjectivity and limiting scalability [7]. The difficulty is compounded in systems characterised by continuous differentiation, where cells occupy transitional states rather than forming discrete clusters, and where the curse of dimensionality can degrade distance metrics and obscure cluster structure [8], [9].

The erythroid lineage offers a well-characterised model of such continuous differentiation, progressing from haematopoietic stem and progenitor cells (HSPCs) through discrete and intermediate stages to mature erythrocytes [10]. Its tightly regulated transcriptional programmes and established marker genes make it an ideal system for studying lineage commitment. Bulk RNA-seq studies, such as Ludwig *et al*. (2019), have provided high-quality reference transcriptomes for distinct erythroid stages [11]. However, translating bulk-derived signatures to the single-cell context is non-trivial due to differences in resolution, technical noise, and the presence of heterogeneous transitional states.

Automated annotation methods, including correlation-based (SingleR [12]), projection-based (scmap [13]), and supervised approaches [14], typically rely on reference atlases or annotated single-cell datasets. While effective in well-studied systems, their performance degrades in specialised biological contexts and finely resolved differentiation hierarchies [7]. Cross-technology generalisation is further limited by systematic distributional differences, even when restricted to transcriptomic measurements from multi-modal assays [15], [16]. Crucially, these methods are not designed to integrate high-confidence bulk-derived signatures with single-cell data, nor to capture the full continuum of erythroid differentiation.

In this study, we address erythroid cell type annotation by com-bining bulk RNA-seq references with expert-curated single-cell data. Starting from the bulk dataset of Ludwig *et al*. (2019) [11], which provides robust but low-resolution cell state definitions, we curate a bone marrow scRNA-seq dataset in which erythroid populations and differentiation stages are manually identified using established biological markers. This expert-annotated dataset captures both canonical and transitional erythroid states at single-cell resolution, bridging bulk-defined identities and single-cell heterogeneity. Using this resource, we train and evaluate supervised classifiers for erythroid lineage annotation and assess their robustness on independent bulk and single-cell datasets, demonstrating that our framework can transfer knowledge across data modalities and reveal finer-grained differentiation structure.

## II. Data & Methods

Our approach consists of two main components (see Figure 1). First, we construct a high-quality, expert-curated reference dataset by integrating a large-scale single-cell bone marrow atlas with biological knowledge derived from bulk RNA-seq of purified erythroid populations. Second, we train supervised classifiers on this curated dataset and evaluate their performance within the original dataset as well as on independent external datasets spanning different data modalities.

**Figure 1.**
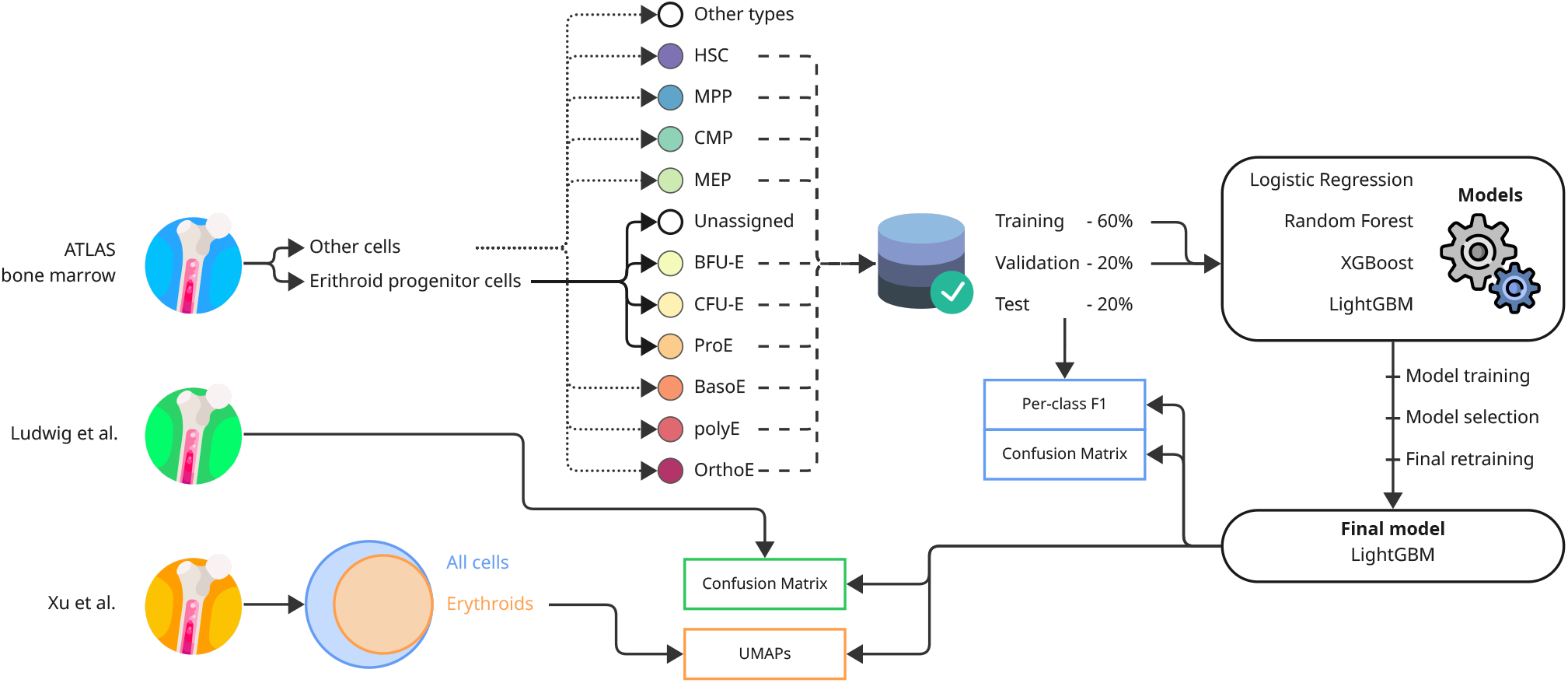
Erythroid lineage classification pipeline. Erythroid cells from three bone marrow scRNA-seq datasets were used to train (ATLAS bone marrow) and benchmark (Ludwig *et al*.; Xu *et al*.) four classifiers (logistic regression, random forest, XGBoost, and LightGBM) along the erythroid differentiation trajectory. The top-performing model (LightGBM) was subsequently retrained on the combined training and validation sets and applied to held-out datasets, with performance evaluated using per-class F1 scores, confusion matrices, and UMAP visualisations.

### A. Construction of an expert-curated erythroid reference dataset

We started from a publicly available scRNA-seq atlas of human bone marrow [17], preprocessed following the original study (quality control, normalisation, dimensionality reduction). From this atlas, we retained cells belonging to the following annotated cell types along the erythroid lineage: haematopoietic stem cell (HSC), haematopoietic multipotent progenitor (MPP), common myeloid progenitor (CMP), megakaryocyte-erythroid progenitor (MEP), erythroid progenitor cells, basophilic erythroblast (BasoE), polychromatophilic erythroblast (PolyE), and orthochromatic erythroblast (OrthoE). However, the atlas does not resolve the early erythroid-committed progenitor stages between MEP and BasoE in the erythroid progenitor cells.

To refine erythroid annotations beyond the granularity available in the atlas, we derived canonical marker gene signatures from the bulk RNA-seq dataset of Ludwig *et al*. [11], which provides high-confidence transcriptional profiles for well-defined stages of erythropoiesis. Guided by these signatures, we subdivided the extracted erythroid progenitor cells into four groups: burst-forming unit– erythroid (BFU-E), colony-forming unit–erythroid (CFU-E), pro-erythroblast (ProE), and an unassigned category. Cells that could not be confidently assigned to any of these stages were discarded. BFU-E, CFU-E, and ProE represent early stages of erythroid commitment not resolved in the original atlas annotations.

These populations were defined using established erythroid markers [10], [11]. BFU-E cells were characterised by high expression of the progenitor marker *CD34* [18], moderate *TFRC* (*CD71*) levels [19], and low expression of mature erythroid markers such as *GYPA* (*CD235a*) [20] and *CD36* [21]. CFU-E cells exhibited reduced *CD34* expression together with increased *TFRC, GYPA*, and *CD36* levels, alongside the emergence of *GATA1* expression [22]. Pro-erythroblasts (ProE) were defined by low *CD34* expression and high levels of *TFRC, GYPA, CD36*, and *GATA1*, reflecting a more advanced stage of erythroid commitment [22].

As shown in Figure 2, this curation produces a more continuous and biologically coherent representation of erythroid differentiation. Compared to the original broad-category labels, the refined annotations capture intermediate states and reveal a clearer progression along the erythroid trajectory, with transitions between early progenitor and committed states becoming more distinguishable. This curated dataset forms the basis for subsequent supervised modelling.

**Figure 2.**
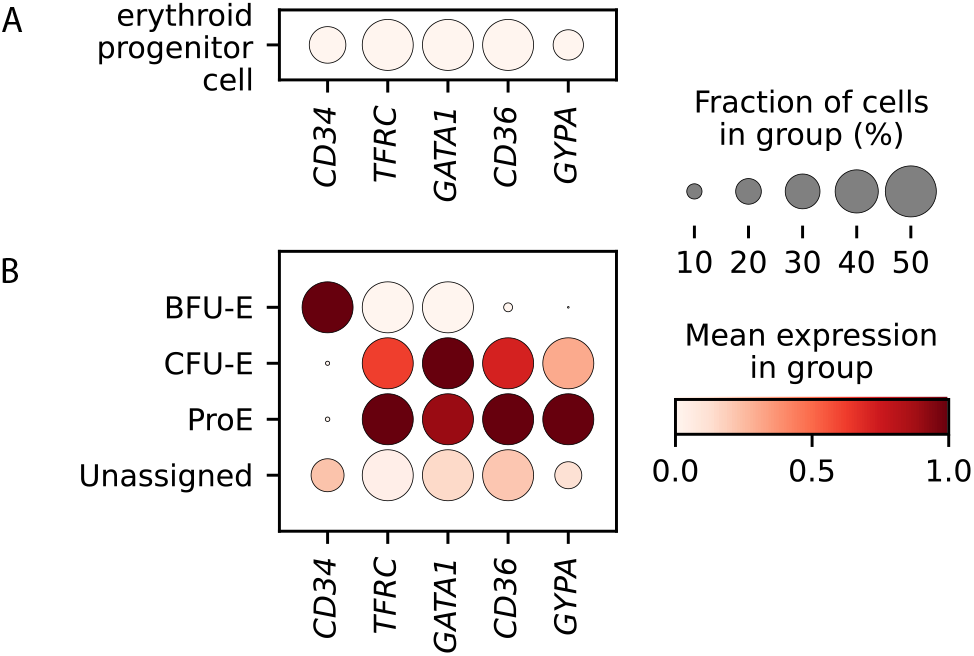
Marker expression in reassigned erythroid progenitors. Dot plots show *CD34, TFRC, GATA1, CD36*, and *GYPA* expression in the original dataset (A) and in subpopulations after reassignment (BFU-E, CFU-E, ProE, unassigned) (B). Dot size indicates the fraction of cells expressing each marker, and colour intensity reflects scaled expression.

The resulting dataset combines the single-cell resolution of the bone marrow atlas with high-confidence stage definitions derived from bulk RNA-seq, yielding a biologically informed training set that captures both canonical erythroid populations and intermediate differentiation states.

### B. Validation datasets

To assess generalisability beyond the training distribution, we employed two independent validation resources.

#### a) Bulk RNA-seq (Ludwig et al.)

The bulk RNA-seq dataset of Ludwig *et al*. (2019) [11] consists of experimentally purified erythroid populations corresponding to well-defined differentiation stages (P1–P8), from early myeloid progenitors to terminal erythroblasts. Bulk RNA-seq reads were preprocessed using CATCH-UP pipeline, in which each sample was processed as an independent replicate [23]. Model predictions were compared to the experimentally defined stage identities, providing a coarse-grained validation of stage-specific signal recovery across data modalities.

#### b) Independent bone marrow scRNA-seq (Xu et al.)

We additionally incorporated a bone marrow dataset from a large-scale cross-dataset harmonisation study [24], comprising approximately 66,000 human cells with harmonised cell type annotations. This dataset integrates single-cell data across technologies, with expert-curated identities organised within a hierarchical framework. For our analysis, we restricted evaluation to cells annotated as belonging to the erythroid lineage, while cells from other lineages were discarded.

Importantly, neither validation dataset was used during model training.

### C. Supervised classification

We formulated cell type identification as a supervised multi-class classification problem. Each cell in the curated single-cell dataset was represented by its projection onto principal components derived from erythroid highly variable genes and associated with a discrete label corresponding to its assigned erythroid differentiation stage.

#### a) Training strategy

The curated dataset was split into training (60%), validation (20%), and test (20%) sets using stratified sampling to preserve class proportions across differentiation stages. This split was performed prior to any preprocessing steps to prevent data leakage. Model training was carried out exclusively on the training subset, the validation set was used for model comparison and selection, and the held-out test set was reserved for final unbiased evaluation. Class imbalance was addressed, where supported, through balanced class weights [25].

#### b) Feature representation and preprocessing

Preprocessing was performed using parameters learned exclusively from the training set. Raw counts were normalised to a fixed library size of 10^4^ counts per cell and log-transformed. Prior to feature selection, genes matching known technical or non-informative categories (including mitochondrial, ribosomal, and non-coding genes) were removed based on standard annotations. Highly variable genes (HVGs; top 2,000) were then identified from the erythroid training cells to reduce dimensionality and mitigate technical noise [6], [26]. Gene expression values were standardised using z-score normalisation, with scaling parameters fitted on the training data and applied to validation and test sets. As an additional dimensionality reduction step, principal component analysis (PCA; 30 components) [27] was applied, with the PCA model fitted exclusively on the training data. To ensure consistency across datasets, genes absent from an evaluation dataset but present in the training feature set were zero-padded to maintain feature alignment across datasets, ensuring that all samples were represented in the same feature space.

#### c) Model selection

We evaluated logistic regression [28], random forests [29], XGBoost [30], and LightGBM [31]. These models were chosen to cover a range of inductive biases: logistic regression serves as a probabilistic baseline, random forests aggregate decision trees through bagging, and XGBoost and LightGBM employ gradient boosting to iteratively combine weak learners. All four are well established for transcriptomic classification tasks [14] and, importantly, permit direct inspection of feature contributions, which could enable downstream biological interpretation of gene-level signals driving classification. While deep learning may offer greater modelling capacity, it typically requires larger training datasets and reduces transparency; given the moderate number of erythroid classes, classical models represent a practical first step. All models were trained with fixed hyperparameter settings and compared on the validation set using macro-averaged F1-score and balanced accuracy. The use of HVGs with PCA-based representations additionally preserves the link between model features and gene expression, enabling downstream interpretability through feature importance analysis and gene loading recovery.

#### d) Evaluation

Within-dataset performance was assessed on the held-out test set using macro-averaged F1-score and balanced accuracy [32], supplemented by confusion matrices for class-level analysis. Cross-dataset evaluation was performed by applying the selected model to the Ludwig *et al*. bulk RNA-seq data and the Xu *et al*. bone marrow scRNA-seq dataset, as described in Section II-B.

#### e) Implementation details

Logistic regression and random forests were implemented using `scikit-learn` [33], while gradient boosting models used the `XGBoost` and `LightGBM` libraries. All transformations were fitted exclusively on the training data, and reproducibility was ensured through fixed random seeds and deterministic model configurations.

## III. Results

### A. Expert curation refines erythroid lineage annotation in single-cell data

We first assessed the impact of expert-driven curation on erythroid annotations in the bone marrow scRNA-seq dataset [17]. While the original atlas captures the major compartments of the erythroid lineage, it applies relatively coarse haematopoietic labels to progenitor populations and does not resolve early committed stages. By integrating bulk RNA-seq profiles with canonical marker gene expression, we refined the annotation of erythroid progenitor cells, enabling the identification of distinct subpopulation stages, including BFU-E, CFU-E, and ProE, based on expert-guided interpretation of marker genes (Figure 3).

**Figure 3.**
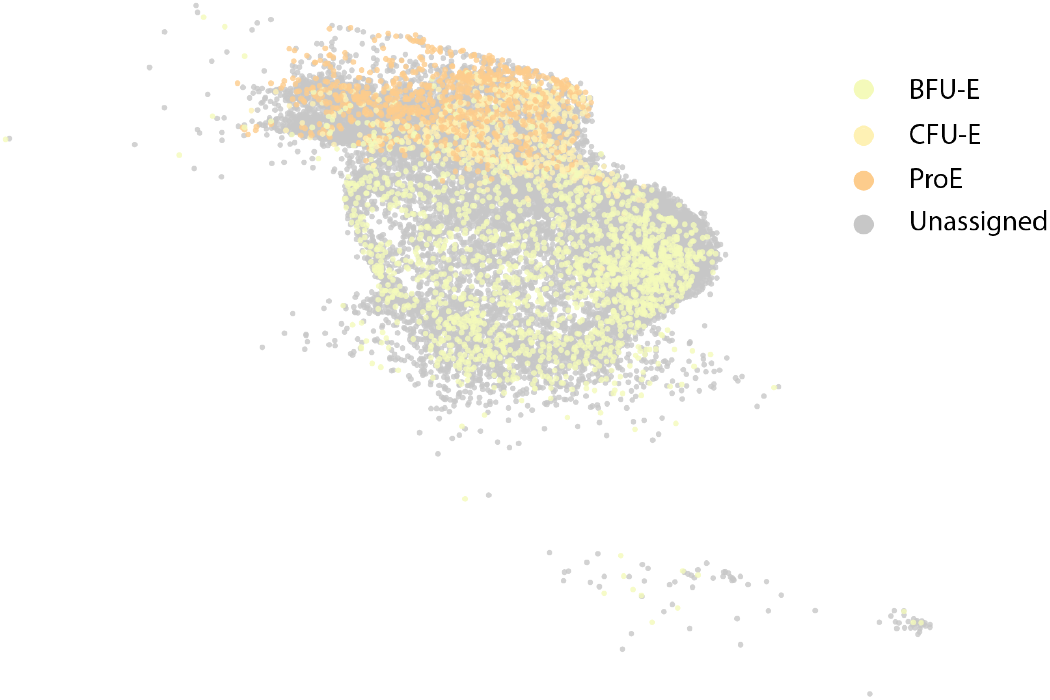
UMAP projection of erythroid progenitor cells scRNA-seq data highlighting refined annotations. Expert-guided curation enabled the identification of distinct erythroid progenitor cells subpopulations (BFU-E, CFU-E, and ProE), based on integration of bulk RNA-seq profiles and canonical marker gene expression. Cells not confidently assigned to a specific erythroid stage are labelled as “Unassigned.” The spatial distribution suggests a continuum of erythroid differentiation from BFU-E through CFU-E to ProE.

### B. Model comparison and selection

We evaluated the four candidate classifiers on the validation set. As shown in Figure 4, gradient boosting methods achieved the highest overall performance. LightGBM attained the best balanced accuracy (0.826) and a competitive macro F1-score (0.821). Based on these results, LightGBM was selected as the final model and retrained on the training and validation sets combined before applying it to the held-out test set.

**Figure 4.**
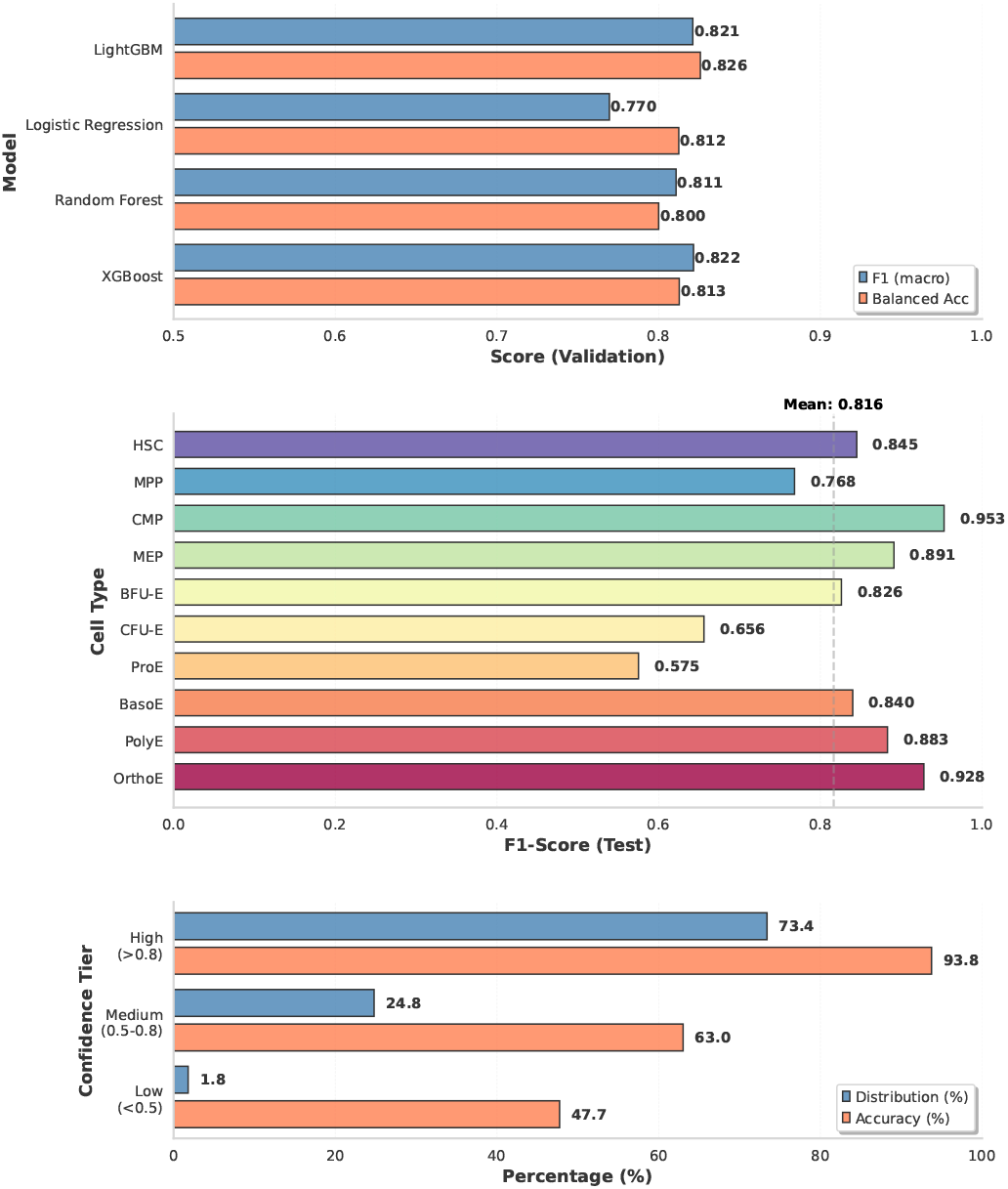
Performance analysis of supervised models for erythroid cell type annotation. (Top) Validation performance of logistic regression, random forests, XGBoost, and LightGBM models, evaluated using macro-averaged F1-score and balanced accuracy. Gradient boosting methods (LightGBM and XGBoost) achieve the highest overall performance, with LightGBM selected for downstream analysis. (Centre) Per-class F1-scores on the held-out test set for each erythroid differentiation stage. Higher performance is observed for more differentiated cell types, while early progenitor populations (e.g., BFU-E, CFU-E) show lower accuracy. (Bottom) Distribution of prediction performance across confidence tiers. The majority of cell types are classified with high confidence (F1 *>* 0.8), indicating robust performance across the erythroid lineage.

### C. Performance on the held-out test set

We evaluated the selected LightGBM model on the held-out test set to obtain an unbiased assessment of classification performance across erythroid stages. Per-class F1-scores (Figure 4, centre) reveal that more differentiated populations are classified with high accuracy: OrthoE (F1 = 0.928), PolyE (F1 = 0.883). Early progenitor states were more challenging, with BFU-E (F1 = 0.656) and CFU-E (F1 = 0.575) showing lower scores, likely reflecting their transitional transcriptional profiles and proximity in gene expression space.

Grouping predictions by confidence tiers, the majority of cell types (73.4%) fall into the high-confidence tier, 24.8% show intermediate performance, and only 1.8% are classified with low confidence (Figure 4, bottom).

The normalised confusion matrix (Figure 5) provides further detail. Late erythroid populations and committed progenitors are classified with high accuracy: CMP (0.94), MEP (0.93), OrthoE (0.90), PolyE (0.87), HSC (0.85), BasoE (0.84), and BFU-E (0.82). Performance is lower for CFU-E (0.70) and ProE (0.60), the two stages that sit at the centre of the early erythroid commitment continuum. In both cases, misclassifications involve immediate neighbours: CFU-E cells are primarily confused with ProE (0.20) and BasoE (0.10), while ProE cells spread into BasoE (0.20) and CFU-E (0.18). A similar pattern is observed at the top of the hierarchy, where HSC and MPP show mutual confusion (HSC → MPP 0.15; MPP → HSC 0.17). These errors are consistently local, confined to adjacent stages rather than distant cell types, indicating that the model preserves the biological ordering of the differentiation hierarchy even where predictions are uncertain.

**Figure 5.**
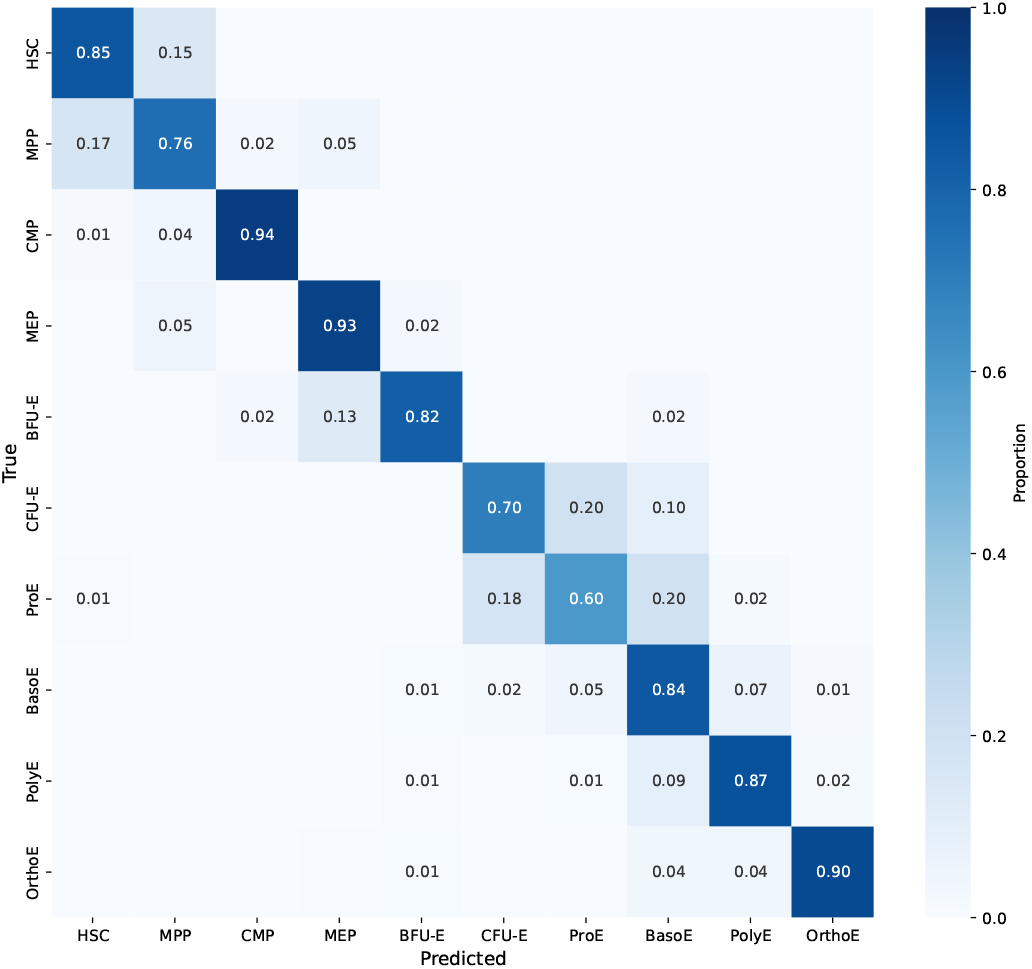
Normalised confusion matrix of erythroid cell type predictions on the held-out test set. Rows correspond to true labels and columns to predicted labels. High diagonal values indicate correct classification. Off-diagonal errors are primarily confined to neighbouring differentiation stages, consistent with gradual transcriptional transitions.

### D. Transfer to bulk erythroid transcriptomes

To assess whether the model captures biologically meaningful erythroid states beyond the single-cell context, we applied the trained LightGBM classifier to bulk RNA-seq samples from Ludwig *et al*. (2019) [11]. For each replicate, we obtained prediction probabilities across erythroid stages. As shown in Figure 6, the model assigns high probabilities to the expected cell types for the majority of samples. Samples enriched for CFU-E (P2) and proerythroblasts (P3–P4) are primarily assigned to intermediate states, while later populations (P5–P8) show strong correspondence with basophilic, polychromatophilic, and OrthoE. Where discrepancies occur, they are limited to neighbouring stages, consistent with gradual transcriptional transitions. Intermediate populations exhibit broader probability distributions across adjacent stages, reflecting the continuous nature of differentiation.

**Figure 6.**
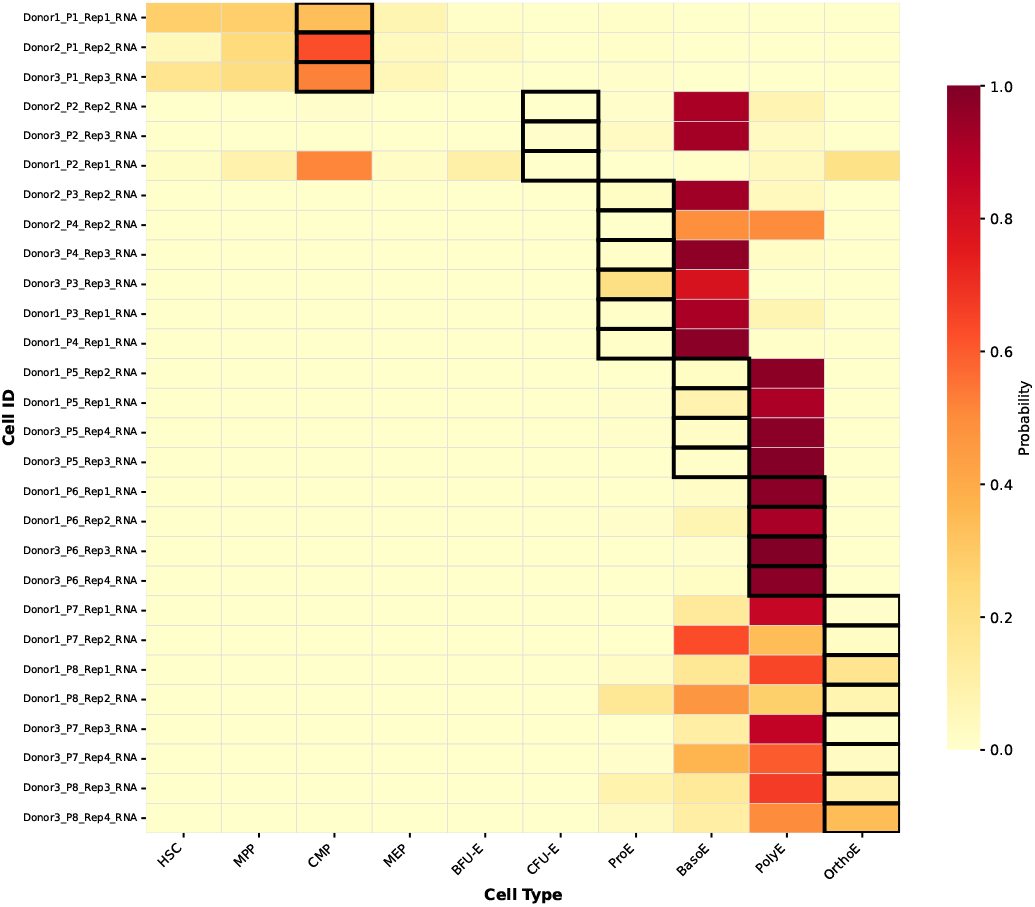
Prediction of erythroid stages in bulk RNA-seq samples from Ludwig *et al*. (2019). Heatmap of predicted probabilities for each erythroid cell type across bulk replicates. Black boxes indicate the expected cell type based on experimental purification. Diagonal enrichment reflects agreement between predicted and expected stages, with deviations confined to neighbouring states.

These results demonstrate that classifiers trained on single-cell data retain sufficient biological signal to recover stage-specific identities in bulk transcriptomic profiles, supporting both the validity of the curated training dataset and indicating that the model can effectively transfer across data modalities.

### E. Refinement of erythroid annotation in an independent single-cell dataset

We applied the trained model to the bone marrow dataset of Xu *et al*. [24] to evaluate its applicability to an independently generated single-cell resource. The original annotations group erythroid cells into broad categories (early, mid, and late erythroid), without resolving intermediate stages such as BFU-E, CFU-E, or ProE (Figure 7A).

**Figure 7.**
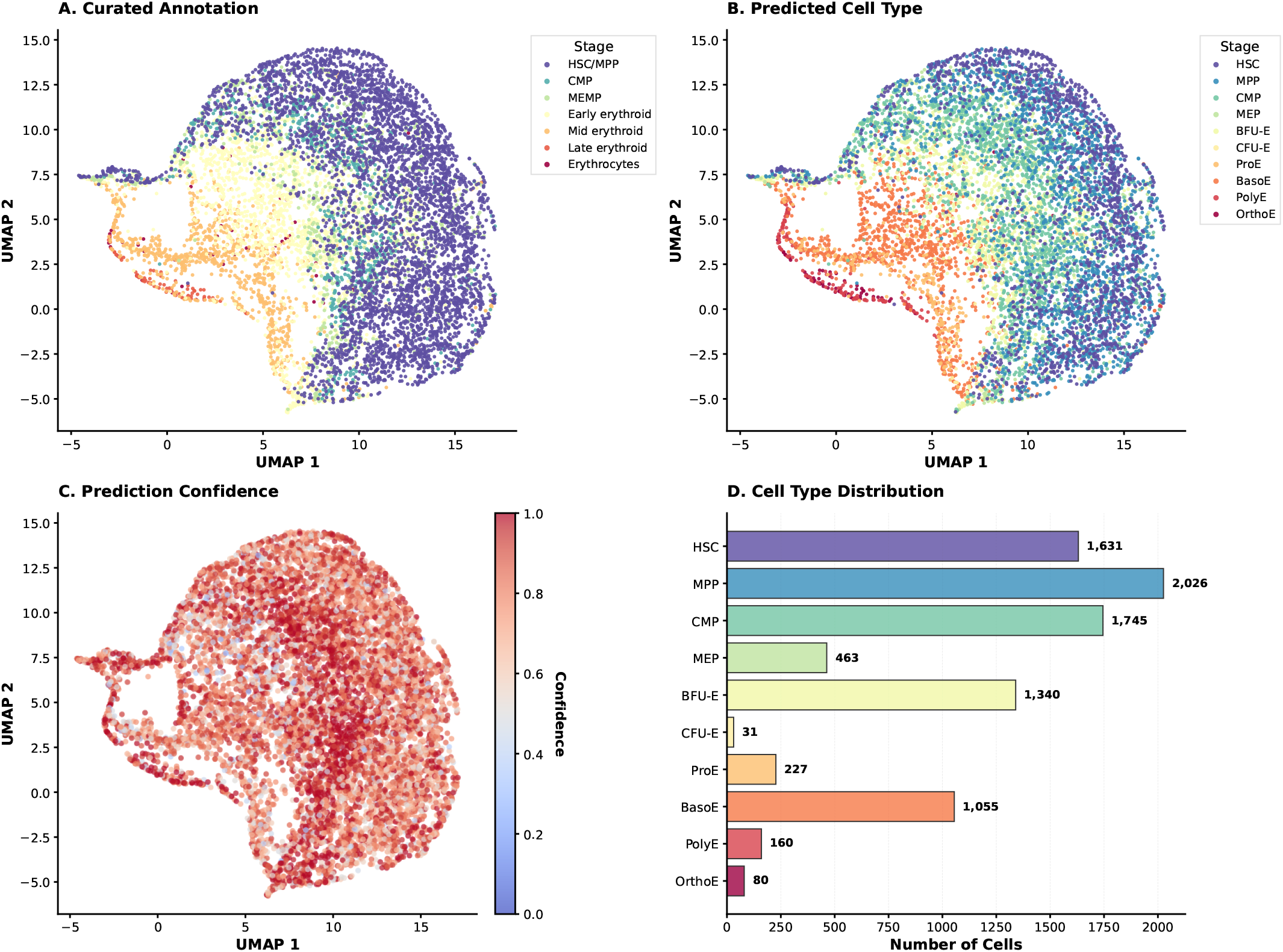
Refinement of erythroid lineage annotation in an independent bone marrow dataset. (**A**) Original annotation showing broad erythroid populations. (**B**) Predicted cell types using the trained LightGBM model, revealing finer-grained differentiation stages. (**C**) Prediction confidence across cells; lower confidence is observed at transitional boundaries between adjacent states. (**D**) Distribution of predicted erythroid cell types, highlighting previously unresolved intermediate populations.

The model predictions (Figure 7B) resolve distinct developmental stages along the differentiation trajectory, further subdividing early progenitors into BFU-E, CFU-E, and ProE, while consistently identifying later stages such as basophilic, polychromatophilic, and OrthoE. The predicted labels exhibit a coherent spatial organisation in the UMAP embedding, delineating a continuous trajectory from stem cells to terminal erythroid states. This organisation is consistent with the continuous nature of erythroid differentiation, where transcriptional changes occur gradually rather than as discrete transitions between cell states, explaining the observed overlap between neighbouring populations. In this context, the model effectively captures intermediate states and gradual transitions in cell identity.

Examination of model confidence (Figure 7C) reveals that the majority of cells are assigned with high certainty, particularly along well-defined trajectory regions. Lower confidence occurs at boundaries between adjacent states, reflecting gradual transcriptional changes. The distribution of predicted cell types (Figure 7D) highlights previously unresolved subpopulations, including substantial numbers of BFU-E and BasoE cells.

These results demonstrate that models trained on expert-curated data can both transfer annotations to new datasets and refine existing labels to reveal finer-grained differentiation structure.

## IV. Conclusion & Future Work

We have presented a supervised framework for erythroid lineage annotation in scRNA-seq data, built on expert-curated labels that integrate bulk and single-cell transcriptomic information. The resulting training dataset resolves intermediate and early progenitor states typically absent from standard annotations.

The approach achieved robust classification across the majority of erythroid stages, with errors predominantly confined to neighbouring states along the differentiation hierarchy. Successful transfer to both bulk RNA-seq and an independent single-cell dataset supports the robustness of the learned representations across data modalities and experimental conditions. In the absence of detailed ground-truth labels in external datasets, expert-driven interpretation provided the primary means of assessment, underscoring both the strengths and current limitations of the approach.

Several challenges remain. The dependence on expert validation highlights the need for more comprehensive and standardised reference datasets, particularly for finely resolved differentiation systems. While HVGs and PCA-based representations enabled effective modelling, systematic investigation of feature importance and gene-level contributions is needed to better understand the biological basis of classification decisions [34]–[Feature attribution methods such as MAGNETO could further refine marker selection and improve interpretability, potentially revealing novel markers for specific erythroid stages [37].

Future extensions include the integration of larger and more diverse training datasets, more advanced dimensionality reduction techniques to capture non-linear expression structure [35], and deep learning approaches that may offer greater modelling capacity at the cost of interpretability. Incorporating multi-omics data, such as single-nucleus and multiome assays, represents a further promising direction for enhancing cell type resolution [38], [39].

Finally, the framework is not limited to erythropoiesis: it may be applied to other differentiation systems where fine-grained annotation is difficult and high-quality training data are scarce. In such contexts, combining expert knowledge with supervised learning offers a practical strategy for improving annotation accuracy and uncovering previously unrecognised cellular states. The effectiveness of such approaches, however, remains contingent on the availability and quality of curated training data, as discussed in [38].

## Acknowledgment

N.S. is supported by the Wellcome Trust grants (225220/Z/22/Z). S.G.R. is supported by the MRC grant (MC UU 00029/3). J.R.H. is supported by the Wellcome Trust grants (225220/Z/22/Z and 106130/Z/14/Z) and the MRC grant (MC UU 00029/3).

## Declaration

J.R.H. is a co-founder and director of Nucleome Therapeutics and provides consultancy to the company.

